# Optimization of psoriasis – like mouse models: A comparative study

**DOI:** 10.1101/2020.03.05.978775

**Authors:** Christina Karamani, Ivi Theodosia Antoniadou, Aikaterini Dimou, Evgenia Andreou, Georgios Kostakis, Asimina Sideri, Andreas Vitsos, Athina Gkavanozi, Ioannis Sfiniadakis, Helen Skaltsa, Georgios Theodoros Papaioannou, Howard Maibach, Michael Rallis

## Abstract

Psoriasis, a common chronic, autoimmune, inflammatory, relapsing disease should benefit from reliable and human relevant animal models in order to pre-clinically test drugs and approach their mechanism of action. Due to ease of use, convenience and low cost, imiquimod (IMQ) induced psoriasis-like model is widely utilized; however, are all mouse strains equivalent, is the hairless mouse utilizable and can the imiquimod model be further optimized? Under similar experimental conditions, common mouse strains (BALB/c, C57BL/6J, ApoE) and a new hairless strain (ApoE/SKH-hr2) as well as several inducers (IMQ, IMQ + Acetic Acid (AcOH) topical and IMQ +AcOH systemic) were compared by clinical, histopathological, biophysical and locomotor activity assessment. Results showed that BALB/c mice yielded optimal psoriasis-like phenotype with IMQ+AcOH topical treatment, C57BL/6J moderate, ApoE mild, while the ApoE/SKH-hr2 mice due to Munro abscess absence in histopathology analysis left doubts about the psoriasis-like acquisition. The locomotor activity of BALB/c mice treated with IMQ, IMQ+AcOH topically and IMQ+AcOH systemically, showed with all treatments, a decreased covered distance and rearing and an increased immobility. In conclusion, BALB/c appears an optimal psoriasis-like model when utilizing IMQ+AcOH topical application.

## 1 INTRODUCTION

Psoriasis is treated by conventional drugs such as methotrexate, cyclosporine, retinoids and dimethyl fumarate and apremilast or by biologics such as anti TNF-α and IL23/IL17 blockers^[1,2,3]^; however, in such treatments there are limitation suggesting the wisdom to develop new interventions^[4,5,6]^.

Innovative drugs typically pass through pre-clinical tests. As psoriasis is a disease dependent on the innate immune system, which cannot be created in vitro, there is no reliable assay in vitro; therefore, a credible psoriasis mimicking animal model is of potential value. Animal models are used to investigate disease pathophysiology, to assess new drugs, their mechanism of action and to evaluate pharmacodynamic/toxicity/pharmacokinetic relationships^[7]^. The most utilized animal is the mouse; for psoriasis 40+ murine models categorized into spontaneous, transgenic, xenograft and induced^[8,9,10]^, provide important knowledge and led to the identification of new drugs; however, none exactly simulates the human situation^[10,11]^. Spontaneous models have limited applications, transgenic mice define only one gene, are expensive and labor consuming, xenograft are technically difficult requiring large quantities of human tissue and the inducible have no specific nature of the obtained inflammation ^[8,9,10]^. The imiquimod (IMQ)-induced model remains the most utilized mouse model due to ease of use, convenience, low cost ^[8]^ and production of acute psoriasis-like skin phenotype, including erythema, scale formation, epidermal thickening, immune cell infiltration and epidermal barrier disruption^[8,9,12]^. Acetic acid has been utilized in the drinking water as a co-factor to IMQ to enhance psoriasis-like signs, due to its immunomodulatory properties^[13,14]^.

Pre-clinically, a new therapeutic strategy for producing psoriatic-like lesions appears the emotional control of mice in association with topical anti-psoriatic treatment. Th17 cells may have a significant role in depression, while high levels of IL17 and IL17A high levels could be involved in psoriasis-associated depression ^[15,16,17]^. In a mouse psoriasis model, the anti-depression and anti-anxiety drug-fluoxetine, significantly contributed to improvement^[18]^

In the IMQ model most widely used strains are C57BL/6J, BALB/c and secondarily, ApoE mice. Each strain has advantages and disadvantages. Are they equivalent? As the above models are hairy, is there a suitable hairless strain? Is the model optimized? Even though a decade passed since van der Fits et al. proposed the model, the above questions remain sub judice^[19]^. Here, using the IMQ psoriasis-like model, under the same experimental conditions, C57BL/6J, BALB/c and ApoE mice were compared and a new strain of hairless mouse included, Apoe/SKH-hr2, resulted from the crossbreeding of SKH-hr2 and ApoE mice. In addition, further optimization effort was made by topically co-administering IMQ with acetic acid in comparison to drinking water dosing^[14]^. Finally, to broaden the psoriasis-like model i.e. to utilize antidepressant/anxiolytic drugs, behavior tests were performed.

## 2 MATERIAL METHODS

### 2.1 Mice

Six to 15 month old, male BALB/c, C57BL/6J, ApoE and ApoE/SKH-hr2 mice were obtained from the animal care facility in the School of Pharmacy, NKUA (EL 25 BIO-06), allowed 7 days to acclimate prior to experiments and housed in an environmentally controlled condition (22-25^°^C, relative humidity 45-55%) with 12h light/dark cycle with free access to food (Nuevo-Farma Efyra, Greece) and water. The experimental procedure was approved by the National Peripheral Veterinary Authority (5768/30-10-2017, 5769/30-10-2017, 05770/30-10-2017, 5620/22-11-2018) after decision of the protocol evaluation committee for animal experimentation. Animal care was performed according to the guidelines established by the European Council Directive 2010/63/EU.

ApoE/SKH-hr2 mice resulted from crossbreeding of ApoE and SKH-hr2 mice after approval of the National Peripheral Veterinary Authority (4044/14-07-2017) and the committee for animal experimentation. ApoE mice is considered as successful model for IMQ induced psoriasis-like model, while other hairless mice are not^[20]^.

### 2.2 Imiquimod based induced psoriasis-like model

Mice were anesthetized by intra-peritoneal injection with a mixture of ketamin (100mg/kg) and xylazine (7 mg/kg). Then an area of about 7.5cm^2^ (3.0 × 2.5 cm) was shaved on the back of each mouse with an electric shaver. This procedure was not followed in the case of ApoE/SKH-hr2 hairless mice. After 24h, mice of each strain were randomly divided in 4 groups (n=5/group). The control group was treated with a daily application of petrolatum, the IMQ group received a daily topical application of 62.5 mg of a commercially available 5%w/w IMQ cream (Modiwart cream, Pharmazac SA, Greece), the IMQ+ AcOH (topical) group received a daily topical application of 62.5 mg of a cream containing 98% commercially available IMQ cream (5%w/w) in which 2%w/w acetic acid (AcOH) was incorporated; The IMQ+ AcOH (per os) group received a daily topical application of 62.5 mg of commercially available IMQ cream (5%w/w) while the drinking water contained 200 mmol/L acetic acid. The experiment was conducted for 6 days.

### 2.3 Scoring psoriasis-like lesions

Psoriasis Area and Severity Index (PASI), used in the clinical psoriasis evaluation^[21]^, which consisted of measurements for erythema, scaling and thickness, was used to monitor and grade severity of the psoriasis-like lesions. Each parameter was scored independently on first and last experiment day (day 7) on a scale from 0-4, where 0= no clinical signs, 1=slight clinical signs, 2=moderate clinical signs, 3= marked clinical signs and 4= very marked clinical signs. Thickening was scored using an electronic digital caliper (Powerfix Profi, Milomex Ltd, Bedfordshire, UK) and erythema using the Mexameter MX18 (Courage + Khazaka Electronic GmbH, Cologne, Germany). Erythema was evaluated from the measured redness increase according to the following scale: 0= 1-60, 1=60-120, 2=120-180, 3=180-240 and 4=240-300 arbitrary redness units.

Cumulative score (erythema plus scaling plus thickening) served as an objective evaluator of the severity of the induced psoriasis (scale 0-12)^[21]^.

### 2.4 Histological analysis

After sacrificing the mice, back skin was removed, fixed in formalin neutral buffered solution (BDH, England) and embedded in paraffin. 5μm sections were cut and stained with haematoxylin and eosin (H&E). The following representative parameters were evaluated in a doubled blinded assessment by an anatomopathologist: inflammatory cell infiltration, parakeratosis, hyperkeratosis, epidermal thickness and Munro abscess. These features were graded on a 0-3 scale, 0=none, 1=slight, 2=moderate, 3=marked and the summary per sample was characterized as the psoriasis histopathologic score (PHS)^[22]^.

### 2.5 Transepidermal water loss (TEWL) and hydration evaluation

TEWL and skin hydration were measured on the first and last experimental day by using a Tewameter® TM 210 (Courage + Khazaka Electronic GmbH, Cologne, Germany) and a Corneometer CM820 (Courage + Khazaka Electronic GmbH, Cologne, Germany) respectively^[23]^.

### 2.6 Locomotor activity test

To investigate move behavior possibly related to anxiety and depression, a locomotor activity test was performed on the BALB/c mice. Here, BALB/c strain was selected as it showed the most extensive psoriasis-like lesions. Male BALB/c mice were placed into individual, transparent, plexiglass chambers and their locomotor activity scored before and 7 days following IMQ induction. Mice were allowed to move freely in the chamber and were videotaped (Nikon D5100 camera, Japan) for 5 minutes. Total distance covered in the chamber and the number of freezing and rearing events were scored offline. Free online Kinovea video analysis software was used for the measurement of the distance covered ^[24]^. Mice were considered to freeze when they stopped moving for at least 3 seconds.

### 2.7. Statistical analysis

Statistical analyses were performed using one-way analysis of variance (ANOVA) followed by LSD post hoc test or student’s T-test where appropriate. Data are presented as means ± SEM. Differences were considered statistically significant at a level of *p<0.05 and **p<0.005. The statistical analyses were performed in IBM SPSS software package (USA).

## 3 RESULTS

### 3.1 Erythema, thickness, scaling, TWEL and hydration changes

All measured parameters in relation to their controls, were statistically significant (p<0.005). Greatest erythema was in ApoE/SKH-hr2 hairless mice, while between treatments, no statistical difference was discerned with the exception of IMQ + AcOH (per os) in the case of ApoE mice (p>0.05, Figure 1a).

**Figure 1.**
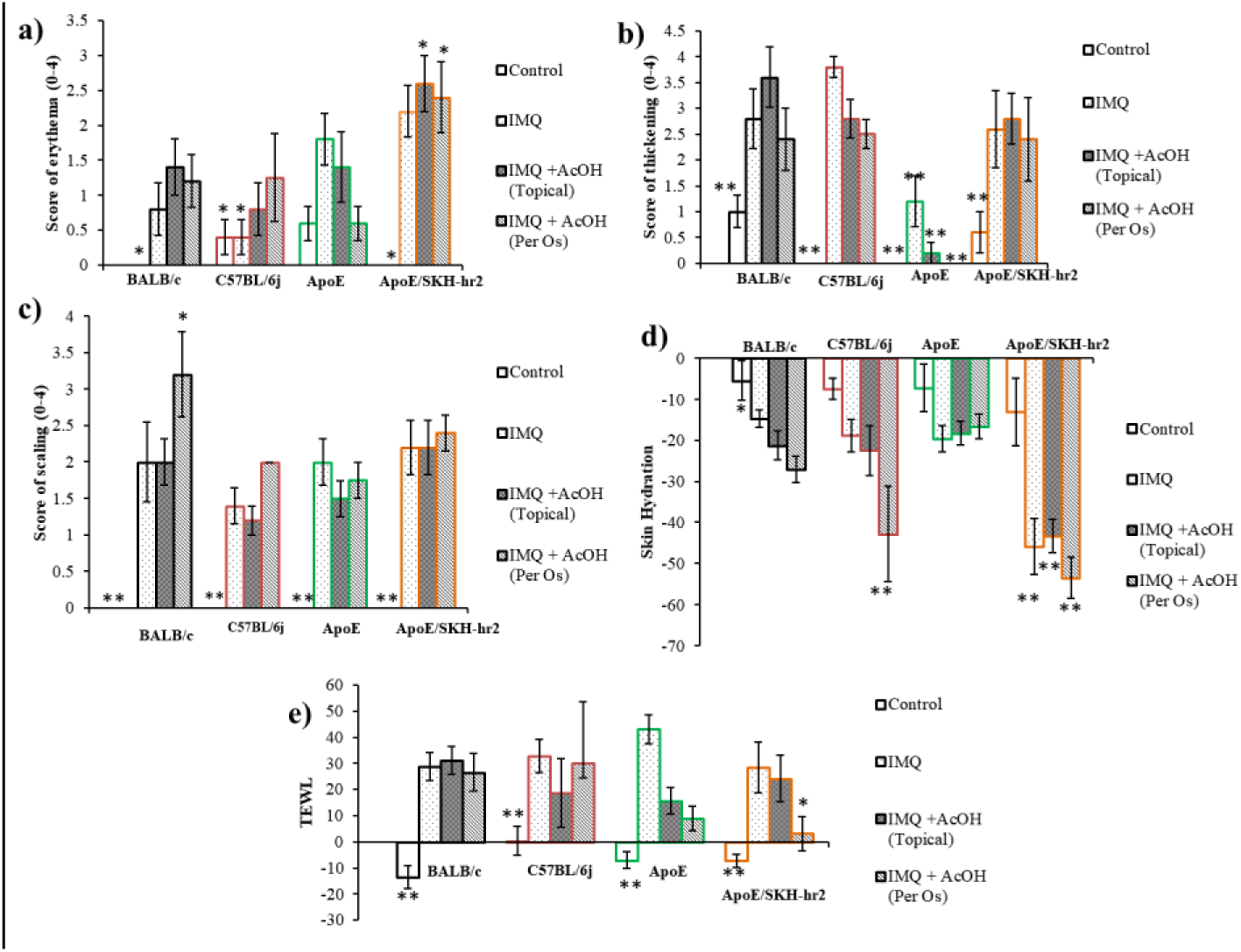
Scores on day 7 of erythema (a), thickness (b) and scaling (c) of psoriasis mouse models(d). Decrease of skin hydration from the first day of the experiment (e). Increase of TEWL from the first day of the experiment. Results are shown as mean ± SEM. *P<0.05, **P<0.005 vs BALB/c mice treated with IMQ + AcOH (topical). *All models showed in relation to their controls statistically significant differences* (p<0.005)

With the exception of ApoE mice, the skin of all other strains was thickened, in average, 2.5 to 3.5 times. In the case of IMQ for C57BL/6J and IMQ + AcOH topically applied for BALB/c mice showed relatively to some of the other cases enhanced thickening (p<0.05, Figure 1b).

With BALB/c and the ApoE/SKH-hr2 mice showed enhanced scaling while IMQ+AcOH (per os) treatment showed the most significant scaling in the case of BALB/c mice (p<0.05, Figure 1c).

The greatest dry skin between all treatments was produced in the ApoE/SKH-hr2 mice (p<0.005). The treatment which caused excess dryness in three of the four strains was IMQ+AcOH (per os) (Figure 1d).

TEWL in all cases was enhanced 3-4 times, demonstrating a significantly decreased barrier function. Greatest TEWL enhancement was obtained by IMQ treatment in ApoE mice (p<0.05, Figure 1e).

### 3.2 PASI scores

Highest mean PASI scores were obtained in ApoE/SKH-hr2 and BALB/c mice. With the exception of ApoE mice, no statistical differences between treatments appeared (p>0.05, figure 2a)

**Figure 2.**
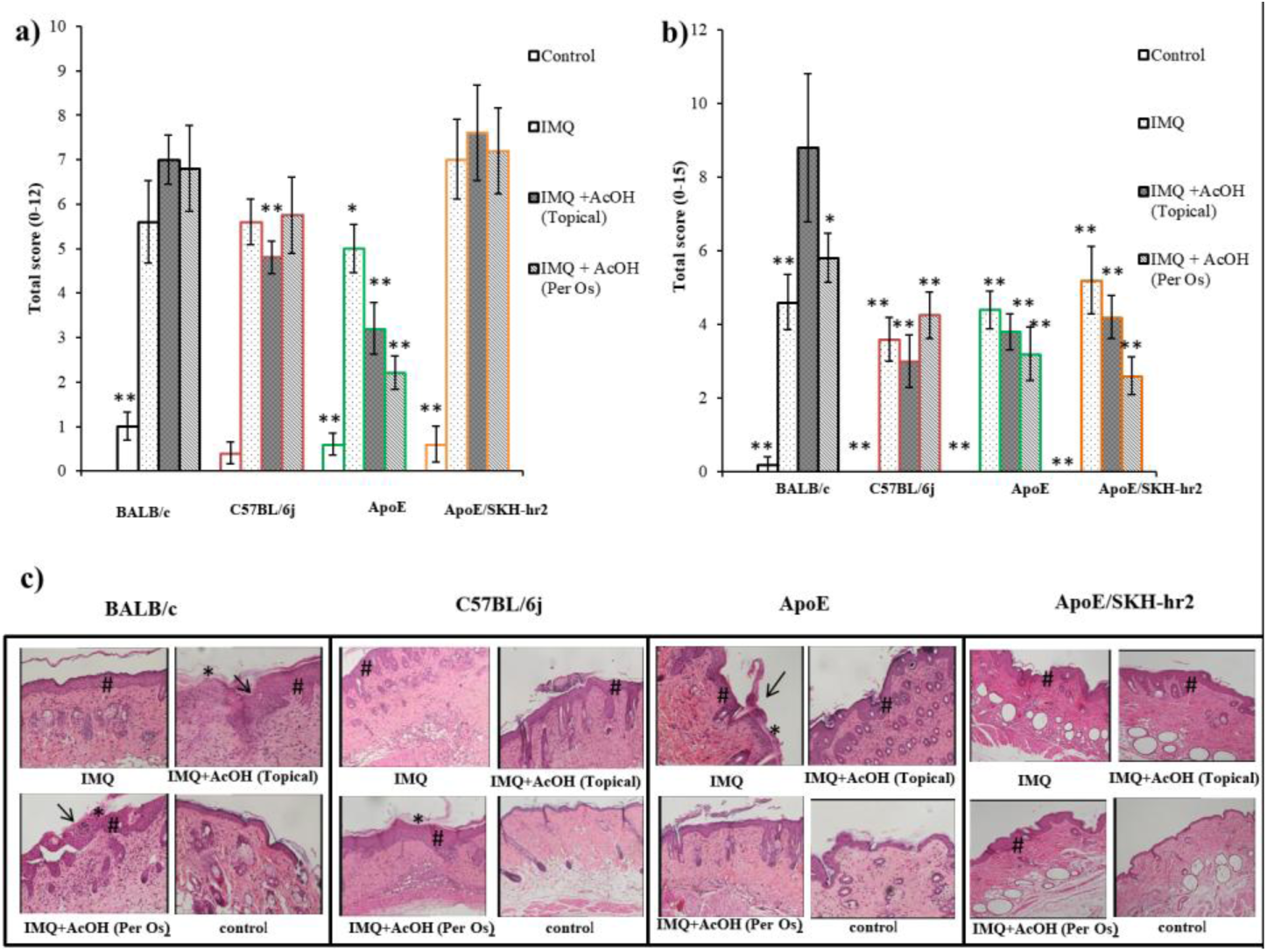
(a) Cumulative PASI score, (b) Psoriasis histopathologic score (PHS), and (c) Histological evaluation of the back mouse skin (H&E staining, magnification 100x) on day 7^th^. Results shown as mean ± SEM. *P<0.05 vs BALB/c mice treated with IMQ+ 2% Acetic acid.*, and 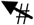 signs indicate parakeratosis, Munros’ microabscess and hyperkeratosis correspondingly. *BALB/c strain and IMQ+AcOH (topical) treatment enhance psoriasis-like phenotype*

### 3.3 PHS score and histopathological findings

Highest statistically significant psoriasis histopathology score (PHS) score was obtained in the BALB/c mice with IMQ+AcOH (topical) treatment (p<0.05, Figure 2b).

Inflammatory cell infiltration, parakeratosis, hyperkeratosis, epidermal thickness and Munro abscess were seen in AcOH treated BALB/c mice. No IMQ sample (n=5) showed Munro abscess, but did with topical application of IMQ+AcOH. According to the histopathology evaluation BALB/c mice treated with IMQ+AcOH showed a typical human psoriasis-like profile, the greatest being with topical application of IMQ+AcOH. For C57BL/6J mice no differences between treatments were observed; scarce Munro abscesses were shown in all cases, giving in total a mediocre psoriasiform reaction. ApoE mice IMQ showed greatest inflammation criteria and few Munro abscess, while the other treatments did not. However total evaluation was negative for ApoE strain as even IMQ gave only a minimal psoriasiform reaction. ApoE/SKH-hr2 mice showed notable inflammatory criteria for all treatments with total Munro abscess absence.

### 3.4 Psoriasis-like morphological features

Photo-documentation of the mice showed the most intense signs in BALB/c followed by ApoE/SKH-hr2, C57BL/6J and ApoE mice strains, while between treatments it was difficult to discern which was the most injurious (Figure 3).

**Figure 3.**
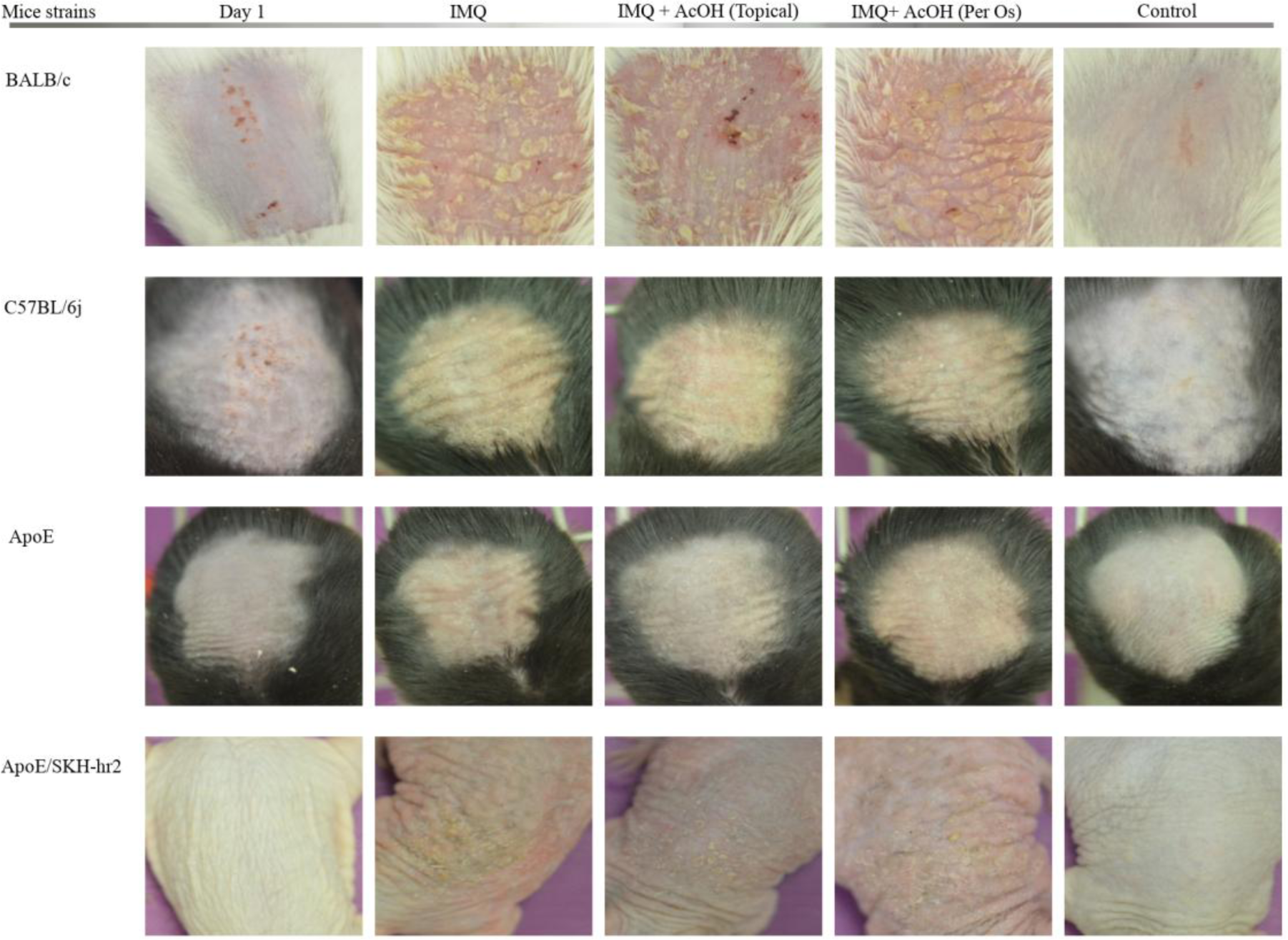
Photodocumetation of mouse back skin on the first and last day of experiment. *BALB/c and ApoE/SKH-hr2 showed the most intense psoriasis-like signs*

### 3.5 Locomotor activity evaluation

Locomotor activity assessment is presented in Figure 4. Treatments significantly decreased the parameters distance covered and rearing after treatment, while the freezing phenomena were in all cases statistically enhanced (p<0.05, Figures 4a, b, c).

**Figure 4.**
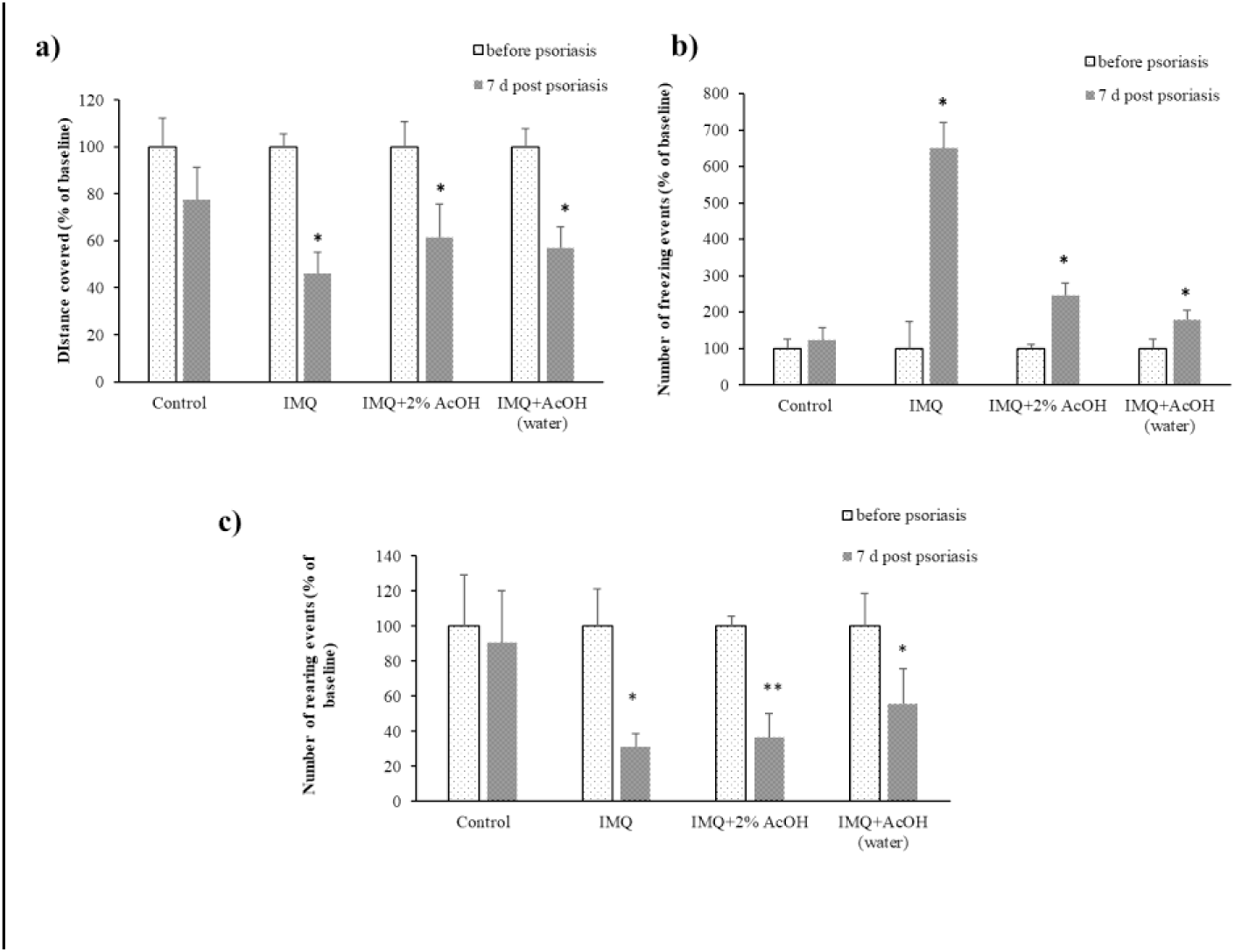
Effect of psoriasis-like induction different treatments on locomotor parameters of BALB/c mice. (A) Distance covered, (B) freezing, (C) rearing were scored before and 7 days following experimental regiments (vehicle, IMQ, IMQ + Topical Acetic Acid, IMQ + Per Os Acetic Acid). Data shown are mean ± SEM. * p<0.05, ** p<0.005 following Student’s t-test. *All interventions showed significant influence on mice behavior* (p<0.05)

## 4 DISCUSSION

Previous studies using hairless mice type SKH-hr1 in which IMQ was topically applied on the back showed no psoriasis-like signs but only dermatitis (unpublished data). This observation accords with that of other investigators stating that “the IMQ model was not induced in this strain’^[20]^. Hairless ApoE/SKH-hr2, (Figure 3) contain melanin and high blood cholesterol values, usually >300 mg/dl, lacking the expression of apolipoprotein E and, therefore, used as an atherosclerosis model ^[27]^. As hypercholesterolemia is associated with psoriasis and use of ApoE mice is described as a psoriasis-like model it is thought that this mouse could be a candidate for a hairless mouse inducing psoriasis-like model ^[26,27]^. Here, in all the clinical and biophysical measured parameters ApoE/SKH-hr2 mice showed significant responsiveness. Erythema, hydration and PASI score gave the higher mean values, statistically different from C57BL/6J and ApoE mice in almost all treatments(Figures 1a, 1c, 2a, p<0.05); however, in its histopathology evaluation showed no Munro abscess, while all the other representative parameters such as cell infiltration, parakeratosis and epidermis hyperplasia were observed. Note that in absence of IL-1 receptor (IL-1R1), or even of IL-17A (IL-17RA) though psoriasis is obtained, in this case, Munro abscess were not created, while in different other psoriasis-like mice models, Munro abscess was obtained at a percentage of 37% ^[28]^. ApoE/SKH-hr2 mice is a new model in which ILs were never measured. This will be helpful in evaluating strain appropriateness of ApoE/SKH-hr2 mouse as a psoriasis model.

Based on our data (Figures 1, 2, 3) and essentially on the most important criteria, principally histopathological evaluation (Figures 2b and 2c), and secondarily, clinical assessment (Figures 2a and 3) the optimized model was the BALB/c strain. Higher PHS values, especially in association with topically applied AcOH (p<0.05), presenting all the significant criteria, inflammatory cell infiltration, parakeratosis, hyperkeratosis, epidermal thickness and Munro abscess; with AcOH treated mice, showing the greatest human like psoriasiform inflammation. Note that immunohistochemical CD4 cells staining analysis, showed that BALB/c strain was more reactive than C57BL/6J or ApoE mice to all treatments (non presented data). C57BL/6J mice showed a relatively close PASI score to BALB/c mice but a small PHS, significantly lower than BALB/c IMQ+AcOH treatments (p<0.005). ApoE mice even though showing a less significant PASI score especially with treatments of IMQ+AcOH (topical) and IMQ+AcOH (per os) histopathological evaluation showed all the criteria less than BALB/c mice for IMQ+AcOH treatments(p<0.005, Figure2a, 2b, 2c). Relating to erythema, scaling, hydration and TEWL, with a few exceptions, the hairy strain mice did not show statistical differences (p>0.05, Figures 2a, 2c, 2d and 2e). For skin thickening, ApoE showed the statistically significant lowest score (p<0.005, Figure 2b). Regarding IL17, IL22 and IL 23 evaluation for C57BL/6J and BALB/c IMQ treated mice, the BALB/c mice have shown higher values in all cases ^[19,29,30,31]^.

The main parameter histopathological evaluation, suggest the optimal treatment, was the topically applied IMQ+AcOH (Figures 2b, 2c). This in the case of BALB/c mice gave higher statistically significant PHS (p<0.05) score with abundant Munro abscesses, while its PASI score was among the highest (Figure 2a, 2b). AcOH per os showed an increase in PHS for BALB/c mice, while the Munro abscesses were seen in the cases of BALB/c and C57BL/6J. IMQ treatment, showed relatively modest result in BALB/c mice, where no Munro abscesses were observed and only ApoE mice had a relatively PASI and PHS significant score obtained in relation to other treatments (p<0.05), in which case, the milder psoriasis-like phenotype is obtained (Figure 2). For PASI score, inflammation, skin thickening, scaling, hydration and TEWL, with some rare exceptions, no statistical differences were observed between treatments (p>0.05, Figures 1, 2a).

Acetic acid topically or systemically administered further optimized the IMQ effect in BALB/c mice. Probiotics administered topically or systemically elicit immune responses on skin, liberating among others, acetic acid^[32]^. Acetic acid significantly influences the immune system, which apparently was a principal reason by which greatest psoriasis-like symptoms were obtained after per os co-administration of acetic acid to topical IMQ in BALB/c mice^[13,14]^. Here on the same BALB/c model showed that the topical co-administration of acetic acid to IMQ further optimizes psoriasis severity (Figure 2). Note that the oral administration of acetic acid at the proposed dose enhanced IMQ toxicity. Mice treated with IMQ lost in mean ∼4% in the case of BALB/c and 8% in the case of ApoE/SKH-hr2 of their weight which doubled in the treatment of IMQ+AcOH (per os) ∼8 and ∼20% correspondingly, while for the topical IMQ+AcOH treatment weight reduction was that of IMQ (non presented data). IMQ treatment weight loss during psoriasis-like induction is confirmed in the literature^[33]^. With IMQ+AcOH (per os) treatment either in C57BL/6J and ApoE/SKH-hr2 mice, one death was noted in each case after excessive weight loss, types of mice with IMQ+AcOH (per os) treatment produced significant reduction of water uptake, which could partially explain the dryer skin obtained with this treatment in three of the four strains (Figure 1d). Previous experiments demonstrated that after ending IMQ treatment, mice quickly recover their previous weight (unpublished data).

IMQ even though topically applied its activity is due to its strong systemic effect influencing thymocytes producing IL17, significant increase of spleen, strongly activating innate immune system^[22,34, 35]^.

In BALB/c strain, which showed the most severe psoriatic-like lesions, locomotor activity of mice was evaluated by distance covered, freezing and rearing. Movement is hypothesized to be related to depression/anxiety symptoms. For that reason different tests based on movements or swim are applied to rodents for behavior evaluation^[36,.37,38]^. Particular emphasis is given to the total distance covered, immobilization time and mice climbing behavior^[37]^. The criteria of total distance, freezing and rearing give a satisfactory appreciation of the mice behavior especially related to depression/anxiety^[37,38]^. Here total distance and rearing significantly decreased and freezing correspondingly increased (p<0.05, Figure 4). Without exception the results showed that the BALB/c psoriasis-like induction by the three treatments significantly influence their behavior as noted (p<0.05). This model may be relevant to humans and could also used to co-evaluate psoriasis-like severity – treatment, as well psychic stress, a parameter so prevalent for psoriasis treatment^[39]^.

Our study adds further questions. Is the topical dose 2%w/w of AcOH optimal or could it be further improved by applying a different dose? The choice of 2% was not arbitrary, as acetic acid is usually used topically in doses ranging from 0.5 to 5%w/w and 2% is frequently applied topically for *pseudomonas aeruginosa* infections^[40,41]^. Another weakness is the non measurability of ILs characteristic of psoriasis like, IL17, IL17A, IL23 and TNF-α in the case of the hairless model ApoE/SKH-hr2. This model, having the advantage of being hairless, is worth further investigation. Note that the greatest PASI score was obtained on the 5^th^ day of application and not on the 7^th^ when the experiment was stopped. Finally, as observed here, we are concerned about mouse weight loss during treatments. Horvath et al, proposed utilizing less IMQ^[33]^. Could this approach be helpful?

In summary, for psoriasis-like IMQ mice models, BALB/c was the most responsive strain. C57BL/6J showed less psoriasis-like lesions, ApoE mild and ApoE/SKH-hr-2 were less useful. The IMQ model can be further optimized by topical co-administration of AcOH while its systematic to IMQ co-administration was toxic. Behavior tests possibly related to depression and anxiety, might broaden the model.

## Abbreviations

PASI: psoriasis area and severity index,
IMQ: imiquimod,
AcOH: acetic acid,
TEWL: transepidermal water loss,
PHS: psoriatic histopathology score,
CD4: T helper cells

## AKNOWLEDGEMENTS

Warm thanks to Mrs MaryAnn Ryder for her valuable contribution in English language, to Mr Giannis Karvelis, Nuevo S.A. company, Greece, for freely providing their high quality mouse chow and Pharmazac SA for kindly providing imiquimod cream (Modiwart).

## AUTHOR CONTRIBUTIONS

MCR, HS, AV, ITA, GTP designed experiments, CK, AD, ITA, EA, GK, AS, AG, IS realized experiments, CK, ITA analyzed data, HIM, MCR wrote manuscript

